# Decoding molecular markers and transcriptional circuitry of naive and primed states of human pluripotency

**DOI:** 10.1101/2020.04.17.046037

**Authors:** Arindam Ghosh, Anup Som

**Affiliations:** Centre of Bioinformatics, Institute of Interdisciplinary Studies University of Allahabad Prayagraj – 211002, INDIA

**Keywords:** Embryonic stem cell, Naive and primed pluripotency, RNA-Seq, WGCNA, Gene module, Hub gene, Enrichment analysis, Transcription factor

## Abstract

Pluripotent stem cells (PSCs) have been observed to occur in two distinct states — naive and primed. Both naive and primed state PSCs can give rise to tissues of all the three germ layers *in vitro* but differ in their potential to generate germline chimera *in vivo.* Understanding the molecular mechanisms that govern these two states of pluripotency in human can open up a plethora of opportunities for studying early embryonic development and in biomedical applications. In this work, we use weighted gene co-expression network (WGCN) approach to identify the key molecular makers and their interactions that define the two distinct pluripotency states. Signed-hybrid WGCN was reconstructed from transcriptomic data (RNA-seq) of naive and primed state pluripotent samples. Our analysis revealed two sets of genes that are involved in establishment and maintenance of naive (4791 genes) and primed (5066 genes) states. The naive state genes were found to be enriched for biological processes and pathways related to metabolic processes while primed state genes were associated with system development. Further, we identified the top 10% genes by intra-modular connectivity as hubs and the hub transcription factors for each group, thus providing a three-tier list of genes associated with naive and primed states of pluripotency in human.

**HIGHLIGHTS:**

- Weighted gene co-expression network analysis (WGCNA) identified 4791 and 5066 genes to be involved in naive and primed states of human pluripotency respectively.
- Functional and pathway enrichment analysis revealed the naive genes were mostly related to metabolic processes and primed genes to system development.
- The top 10% genes based on intra-modular connectivity from each group were defined as hubs.
- Identified 52 and 33 transcription factors among the naive and primed module hubs respectively.
- The transcription factors might play a switch on-off mechanism in induction of the two pluripotent states.

## 1. INTRODUCTION

Conventional human embryonic stem cells (hESCs) derived from the pre-implantation embryo exist in a state quite different from the pre-implantation embryo derived mouse embryonic stem cells (mESCs). It has in fact been found to resemble the post-implantation embryo derived mESCs. Based on these two distinct types of mESCs, pluripotent stem cells (PSCs) have been classified to occur in two states — naive and primed [1,2]. The naive pluripotent cells are assumed to be completely unrestricted and free from any epigenetic constraints. On the other hand, primed pluripotent cells are developmentally advanced and have lineage bias during differentiation [3,4]. This often limits the application of conventional hESCs for biomedical applications [5]. Besides this, the ease of culture of naive PSCs as single cells provides an edge over primed PSCs for genetic manipulation by transfection [6]. These apparent advantages of naive ESCs led to the quest for identification of methods for extraction and/or culture of hESCs in the naive state. Over the last decade, several research groups reported having successfully captured the naive state hESCs (reviewed in [2,3,5,6]). Despite these efforts, the exact differences in molecular mechanisms between the two states still require investigation. Together, naive and primed state pluripotent stem cells represent valuable models to study stages of human development.

Initial efforts to identify transcriptomic markers of the two distinct pluripotency states relied on testing the differential gene expression [7–9]. However, this approach has a critical drawback as it compares each gene in isolation and does not consider the interaction between them [10]. Network-based approach of transcriptomic data analysis has shown to alleviate this problem and has been successful in identifying molecular markers in complex biological systems [11,12]. Co-expression networks, in particular, have been found to be useful in providing a global overview of the co-expression relationships between genes and enables analysis of gene sets [13]. In this work, we use weighted gene co-expression network analysis (WGCNA) to identify the molecular markers that define the two pluripotency states. From our analysis, we identified two clusters of genes (modules) consisting of 4791 and 5066 genes to be associated with naive and primed pluripotent states respectively. Functional and pathway enrichment analysis revealed the naive genes were mostly related to metabolic processes and primed genes to system development. We refined this list to identify the genes with the highest connectivity (hubs) and further the hub transcription factors (TFs) that regulate the expression of these genes. Thus, we provide a three-tier list of genes and their networks that may be crucial to understand the underlying mechanisms that define the two distinct pluripotency states in human. We anticipate that our current findings will lay the groundwork for future experimental studies to understand the underlying mechanisms that define naive and primed pluripotent states.

## 2. MATERIALS AND METHODS

### 2.1. Data retrieval

We used five datasets containing transcriptome profiles of 23 naive and 15 primed state human embryonic stem cell samples in our study (Table 1). The included naive samples were generated by different methods including those by forced expression of pluripotency TFs [14], by the use of small molecule and/or growth factors in culture medium [7,15,16] and direct derivation from an embryo [17]. The datasets were identified based on the criteria that each dataset contained both naive and primed state samples. In addition to this, only those datasets were included in which the library selection for sequencing was done by either poly-A selection or rRNA depletion. The FASTQ files of the 38 samples were retrieved from the European Nucleotide Archive (ENA) [18] and used for downstream analysis. Details of the samples used in this study can be found in Supplementary File 1.

**Table 1:** Summary of data included in the study.

| Project ID | Number of naive samples | Number of primed samples | Reference |
| --- | --- | --- | --- |
| PRJNA286204 | 2 | 2 | [7] |
| PRJEB7132 | 3 | 3 | [14] |
| PRJNA360413 | 3 | 3 | [15] |
| PRJNA309166 | 6 | 4 | [16] |
| PRJEB20388 | 9 | 3 | [17] |

### 2.2. Data processing

The quality of the raw reads was assessed using FastQC v0.11.5 toolkit [19]. Adapters and low-quality reads were trimmed using Trimmomatic v0.36 [20]. HiSat2 v2.1.0 [21] was then used for mapping the raw reads to the reference genome GRCh38.p5 (Ensembl release 84). The expression for each gene was evaluated using featureCounts v1.6.2 [22] based on annotation from Ensembl release 84. Till this stage, each dataset was handled independently. The gene counts from the five datasets were then merged and the low expressed genes which did not have more than one count per million reads (1 CPM) in at least fifty percent of the samples were removed from further analysis. Normalization of the raw reads was performed using DESeq2 and the function removeBatchEffect() from the R package limma was used to account for the different batches [23,24]. Clustering of the samples were performed on the variance stabilized gene counts using hclust() function in R.

### 2.3. Weighted gene co-expression network reconstruction

Reconstruction of the weighted gene co-expression network was done using the R package *“WGCNA”* [25]. The network was constructed by first calculating the Pearson’s correlation coefficient between all possible gene pairs and then raising it to a power β (soft-threshold) to create the adjacency matrix. The soft-threshold is chosen based on the criteria of approximating the scale-free topology of the network [26]. This is followed by generation of the Topological Overlap Measure (TOM) matrix and average linkage hierarchical clustering based on the TOM to clusters genes and identify modules (groups of densely interconnected genes). In our analysis, we identified β=12, corresponding to a scale-free topology fit index R^2^=0.9 using the pickSoftThreshold function in WGCNA (Figure 1). The blockwiseModules function, with deepSplit=2 and minimumModuleSize=30, was used for network construction and module detection. Each module was assigned a unique colour for easy reference in downstream analyses. To determine whether the identified modules were significantly associated with either of the two sample types, a binary matrix was generated describing the association of the samples with their respective traits (naive=0 and primed=1). This matrix was then used as an input to calculate the correlation and p-value with naive and primed sample types.

**Figure 1:**
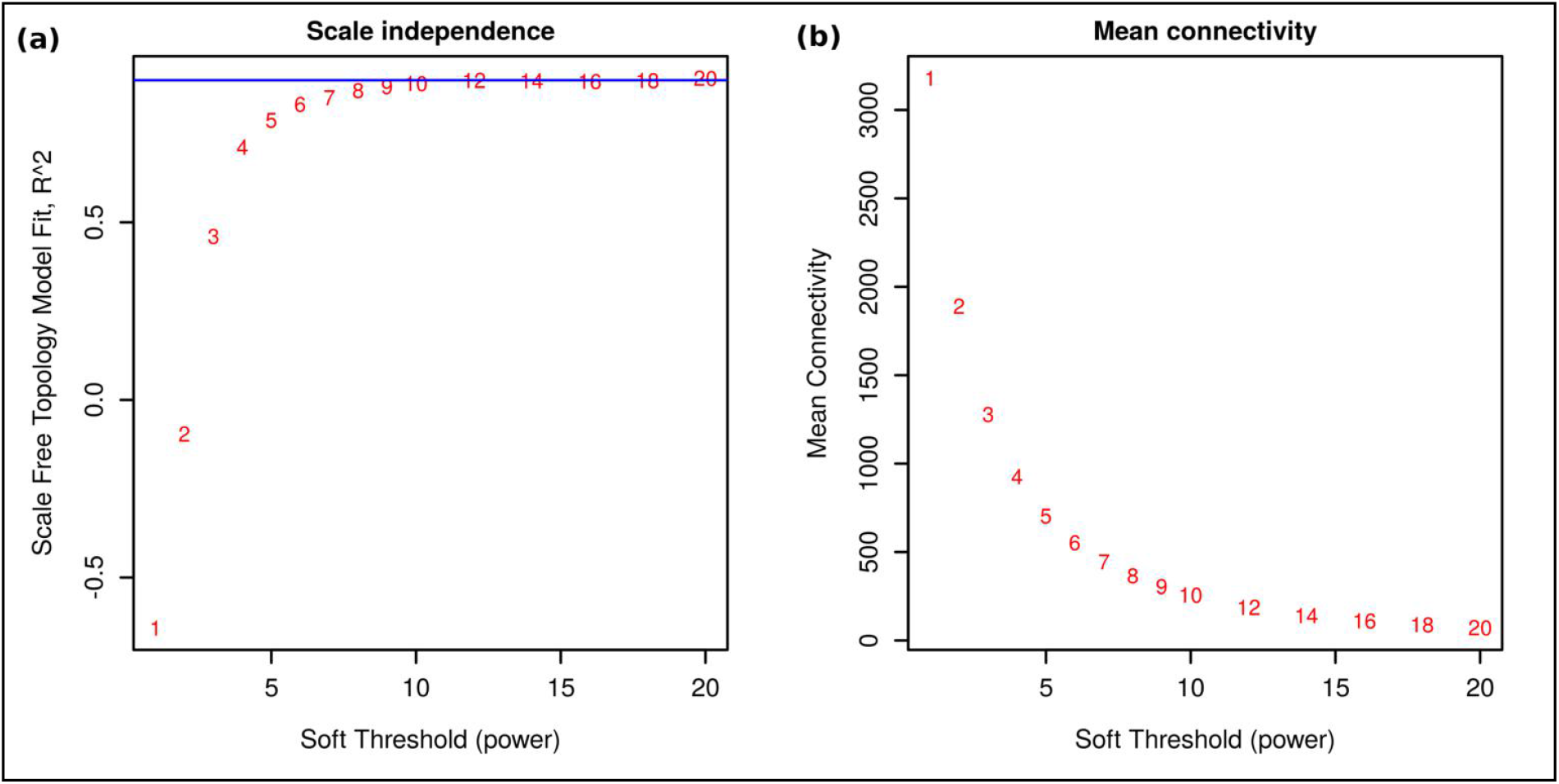
Determination of soft–thresholding power in WGCNA.

### 2.4. Functional and pathway enrichment analysis

Identification of the enriched biological processes and pathways was carried out using PANTHER (Protein ANalysis THrough Evolutionary Relationships) [27]. For biological processes, the annotations from Gene Ontology (GO) database (Released 2020-01-03) and for pathways, Reactome version (65 Released 2019-03-12) were used [28]. Significance testing of the matched terms was performed using Fisher’s exact test. Significantly enriched terms were selected based on false discovery rate (FDR) adjusted p-value < 0.05. REVIGO was used for summarizing and visualization of the gene ontology terms [29].

## 3. RESULTS AND DISCUSSIONS

### 3.1. Clustering of samples

Prior to using the gene count for downstream analysis, we verified whether the samples grouped according to the distinct pluripotency states based on their gene expression. Based on hierarchical clustering and sample-to-sample distance heatmap, no clustering was observed when log-transformed raw gene counts were used (Figure 2a). However, after normalization and batch effect correction, the samples grouped based on their phenotype (Figure 2b). Thus, despite the differences in methods of extraction and culture of the naive/primed state cells they show similar expression profiles. The experimental differences during measurement of gene expression might initially shadow the phenotype-specific differences but once it has been accounted for, the differences due to phenotype are clearly visible. This also sets a way for combining data from multiple RNA-seq experiments to gain statistical power for downstream analysis.

**Figure 2:**
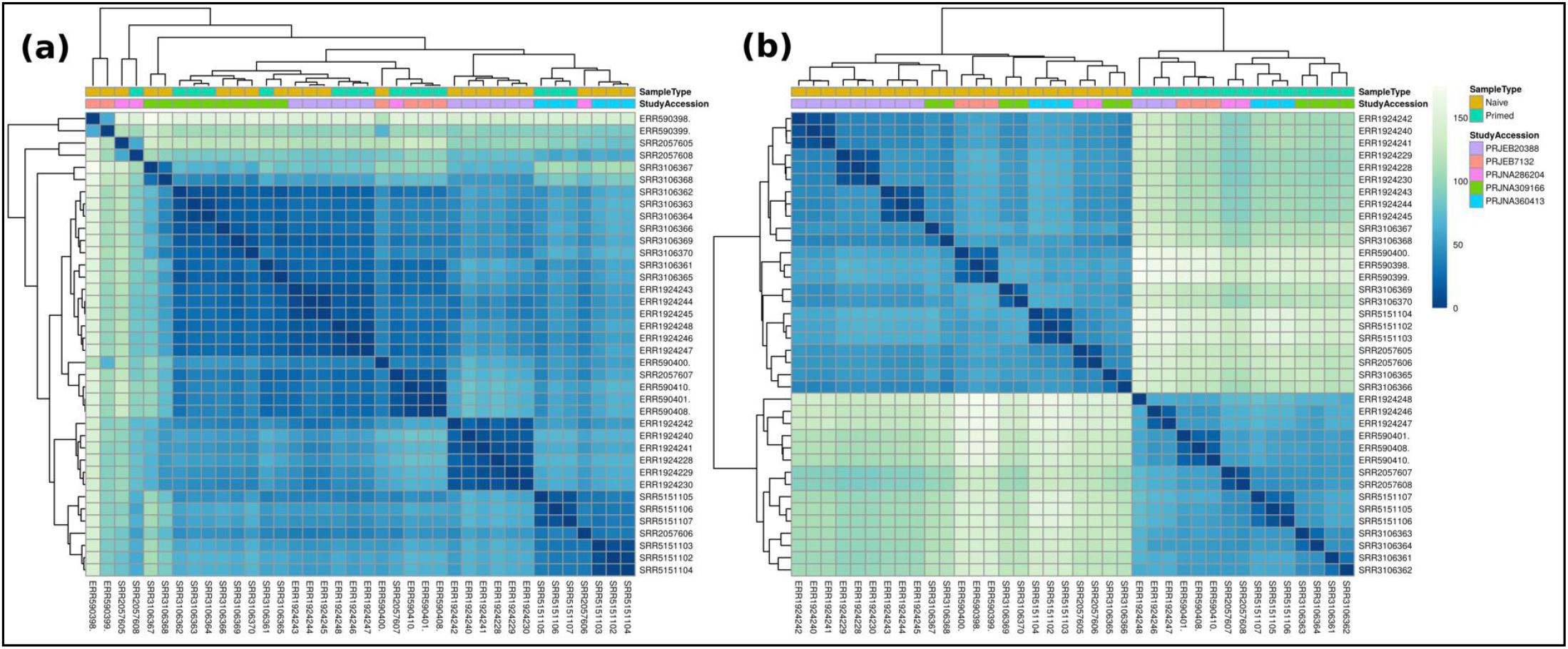
Hierarchical clustering of the samples (a) before and (b) after normalization of gene expression.

### 3.2. Network reconstruction and identification of gene modules

Methods for gene selection based on differential expression analysis only consider individual features in isolation and do not take into account their interaction [10]. Network-based approach has shown to alleviate this problem and helps in the identification of more reproducible marker genes. We used WGCN approach to study the interactions between the entire set of protein-coding genes and identify potential pluripotency state-specific markers. WGCNs are of two major types — signed and unsigned, depending on how the adjacency matrix is calculated. In the signed version, the raw correlation coefficient value is used while the unsigned network uses the absolute value of the correlation coefficient ignoring the direction of correlation. A previous work on mESCs had shown that signed networks provide better system-level understanding of the regulatory mechanisms of ES cells than an unsigned network [30]. Hence, in this work we employed signed hybrid network, a variant of the signed network, to study the patters of co-expression and potential interaction between the genes.

Using the WGCNA approach, we identified 10 modules of highly interconnected genes (Figure 3a). Each of these modules, identified by a unique colour, contained between 60 to 5066 genes (Figure 3b). The genes that could not be assigned into any of the 10 modules are represented within a separate grey module. These modules are representative of a group of genes that work in a coordinated manner for performing a specific biological function. The cumulative gene expression of all the genes in a module is represented by a module eigengene which is essentially the first principal component of the module. The module eigengenes were used in performing module-trait relationship analysis to identify the modules that might be associated with either of the PSC types in our study. Two modules — turquoise (cor = 0.96, p-val = 2e-22) and blue (cor = −0.94, p-val = 9e-19) were found be most significantly correlated to the sample type (Figure 4) and hence were considered for further analysis. Heatmap of the expression profiles of the two significant module genes revealed the blue module genes (4791 genes) showed higher expression in the naive state while turquoise module genes (5066 genes) to have high expression in the primed state (Figure 5). Thus we hypothesised that the blue module genes are critical to the naive state of pluripotency while those of turquoise module are critical to the primed state of pluripotency. The list of genes in the two modules have been provided in Supplementary File 2.

**Figure 3:**
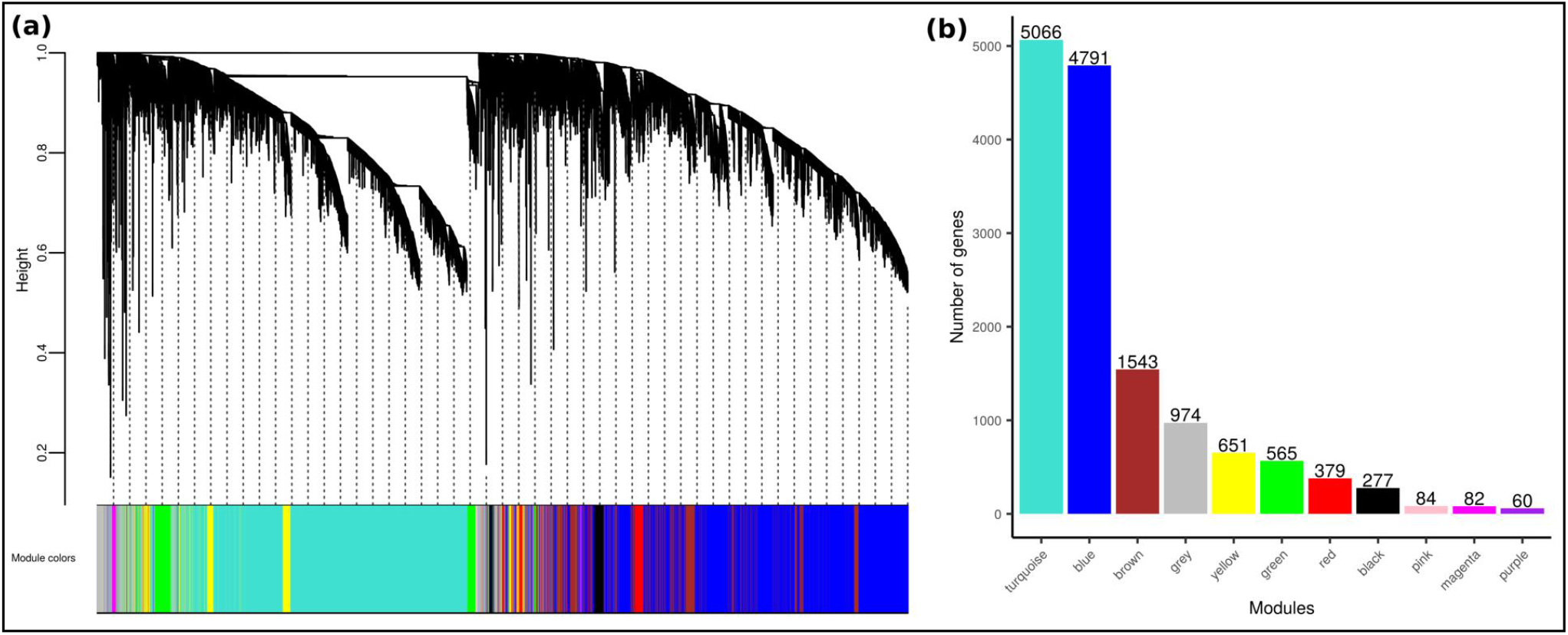
(a) Hierarchical clustering tree of the entire set of protein-coding genes. (b) Bar–plot depicting the number of genes in each module.

**Figure 4:**
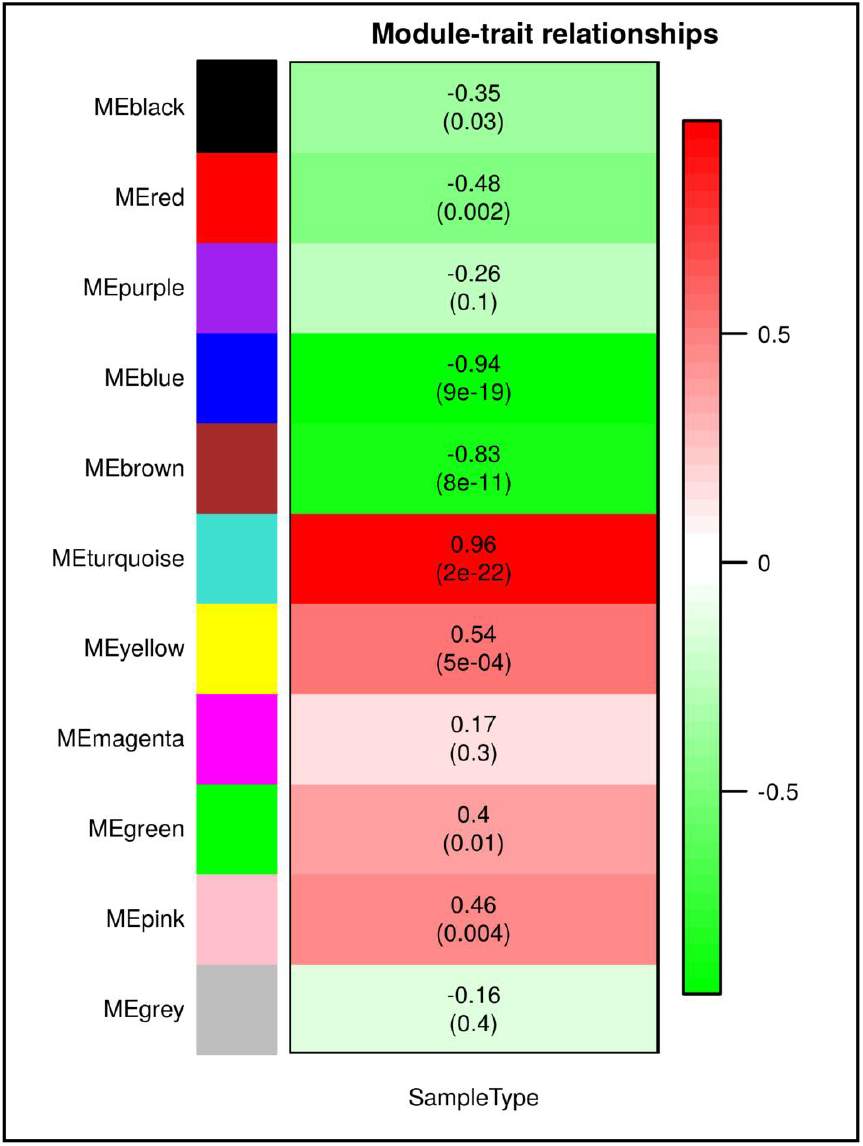
Module–trait relationship analysis. The values within each cell represent the correlation coefficient of the module–eigen with sample type (trait). The corresponding p–values for each module are mentioned within parentheses.

**Figure 5:**
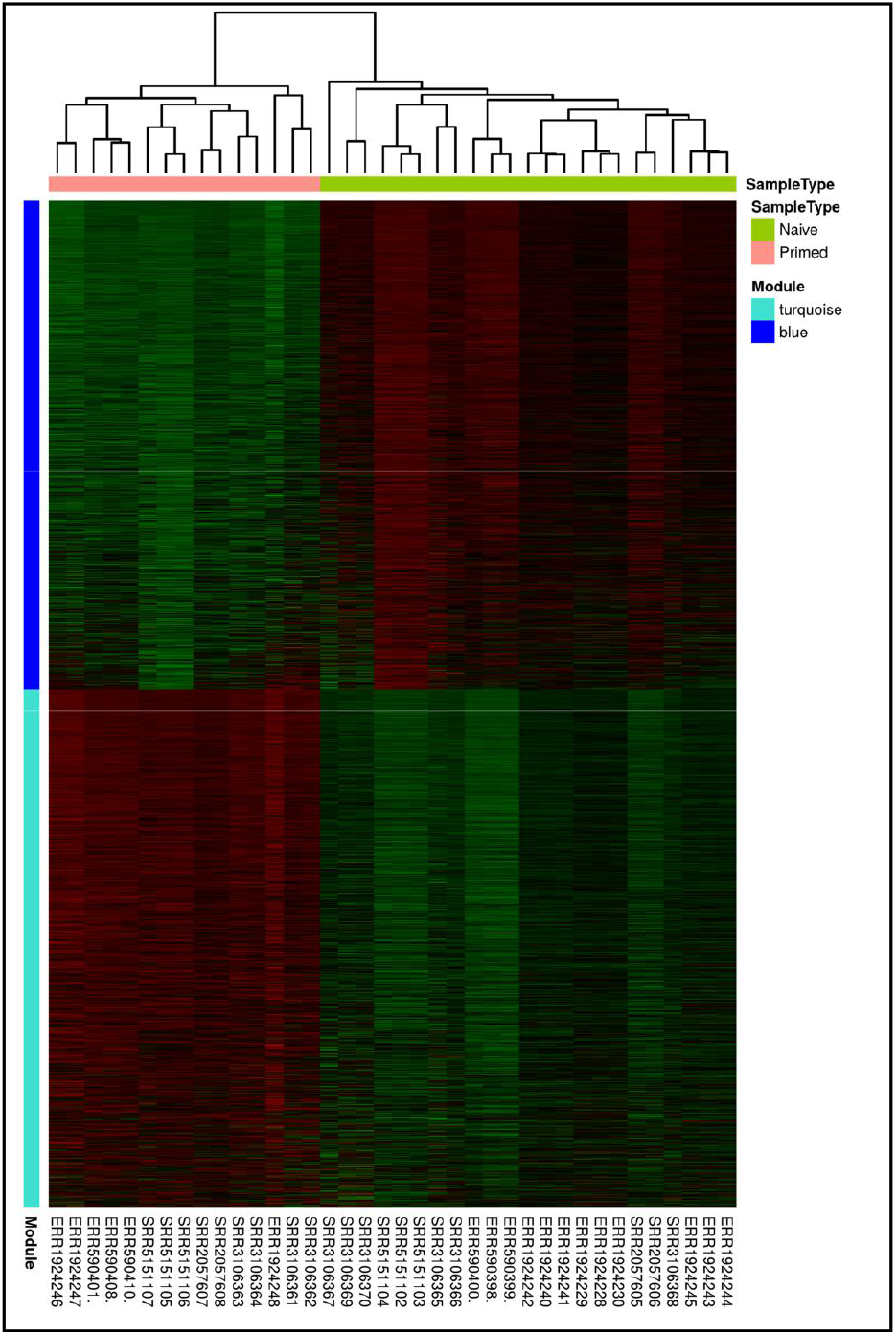
Heatmap of expression of blue (naive) and turquoise (primed) module genes. Here, rows corresponding to genes and columns corresponding to samples. The colour red indicates increased expression, black neutral expression, green decreased expression.

### 3.3. Functional and pathway enrichment analysis of the significant modules

Enrichment analysis of biological processes terms was carried to gain insights into the distinct functions of the genes in the two significant modules. We observed the blue module genes to be mostly involved in metabolic processes like tRNA metabolism, heterocycle metabolism, post-transcriptional regulation of gene expression, cellular respiration etc. (Figure 6a). We also observed terms related to cellular response to DNA damage stimulus enriched in this module. For cells that occur very early in the course of development, maintaining genomic integrity is of utmost importance else DNA damage may compromise the daughter cells. Thus, this explains the enrichment of terms related to metabolism and response to stress in naive pluripotent cells [31]. Besides these, the terms related to mitochondrial biogenesis can also be observed. This is in concordance with previous reports that mitochondrial metabolism is a functional marker of pluripotency stage. Naive PSCs are bivalent in their energy production and can dynamically switch from glycolysis to mitochondrial respiration on demand [32]. In contrast, primed PSCs depend only on glycolysis for its energy requirement. The turquoise module genes, on the other hand, were associated with terms related to system development like regulation of neurogenesis, heart development, eye development, gland development etc. (Figure 6b). This might be due to the fact that primed stem cells are more developmentally advanced than their naive counterpart. Also, it has been found that loss of pluripotency occurs during the transition from naive to the primed state of pluripotency [33]. Similarly, we found the pathways related to tRNA processing, metabolism of RNA, translation, and mitochondrial biogenesis among the significantly enriched pathways in the blue module. The turquoise module genes showed an enrichment of pathways related to extracellular matrix organization, neurotransmitter receptors and post-synaptic signal transmission, laminin interactions etc (Supplementary File 3). Thus, the results of the functional and pathway analysis confirm the hypothesis that the blue module genes are critical to the naive state of pluripotency while those of turquoise module are critical to the primed state of pluripotency.

**Figure 6:**
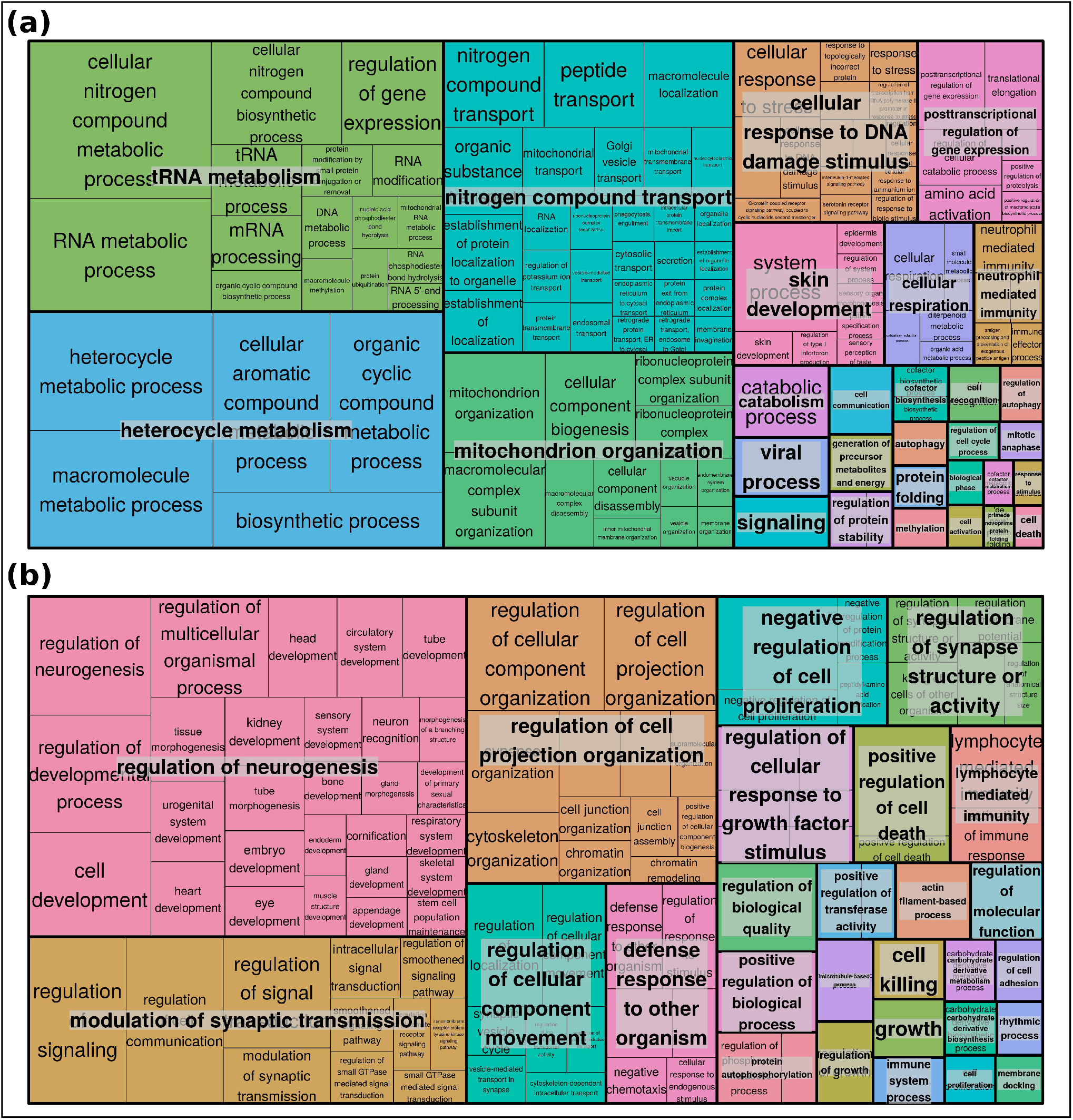
REVIGO treemap representing the most significantly enriched unique GO biological processes terms associated with (a) blue and (b) turquoise modules. Here, colours indicate GO families in which the gene fall and larger boxes indicate a smaller p–value.

### 3.4. Identification of intra-modular hub genes

The nodes with high connectivity (that greatly exceeds the average connectivity) in a network are called “hubs” and are thought to serve specific purposes in their network. Hubs have a significant impact on the network structure and function — hubs serve as bridges between the small degree nodes (i.e., shorting the path lengths in a network) allowing effective/quick flow of information within the network components, and are responsible for network robustness and attack tolerance [34]. The intra-modular connectivity of the genes in the blue and turquoise modules was calculated and the top 10% genes (479 genes in blue and 507 genes in turquoise) with highest intra-modular connectivity were considered as hubs (Supplementary File 4). The network of the hub genes in blue and turquoise modules connected by the top 1% edges by weight within each module are shown in Figure 7. These hub genes included several well established naive and primed state pluripotency markers. For example, the blue module hubs include naive state specific genes like ALPP, ALPPL2, ASH2L, DNMT3L, DPPA3, DPPA5, FAM151A, GDF3, HORMAD1, HYAL4, KHDC3L, OLAH, SMARCA2, SUSD2, and TRIM60. On the other hand, the primed state specific genes ALCAM, CD24, CDH2, FAT3, HLA-A, JMJD1C, KLHL4, NOTCH2, NOTCH3, PTPRZ1, and SIRPA were identified as turquoise module hub genes [5]. Among these, ALPPL2 and SUSD2 are naive state specific surface markers [35,36] while CD24 and HLA-A are primed state specific surface markers [37] and are routinely used for monitoring the primed-to-naive transition of hESCs in culture. DNMT3L is a key regulator of de novo DNA methylation [38]. DPPA3 is a gene critical for Lin28a-regulated conversion from naive to primed pluripotent state [39]. GDF3 helps in activating the Nodal signalling pathway and helps in regulation of hESCs’ ability to maintain the undifferentiated state [40]. KHDC3L is responsible for robust DNA damage response and enhanced homologue recombination mediated repair [41]. The primed state markers ALCAM and CDH2 are cell adhesion proteins and promote homotypic cell-cell contacts. A study by Takehara et al [42] found that CDH2 is important for the maintenance of mouse epiblast stem cell in an undifferentiated state. JMJD1C is a histone demethylation protein that is known to repress neural differentiation of hESCs and hence maintaining its identity [43]. NOTCH2 and NOTCH3 are receptors in the Notch signalling pathway and are involved in cell fate determination during embryonic development [44].

**Figure 7:**
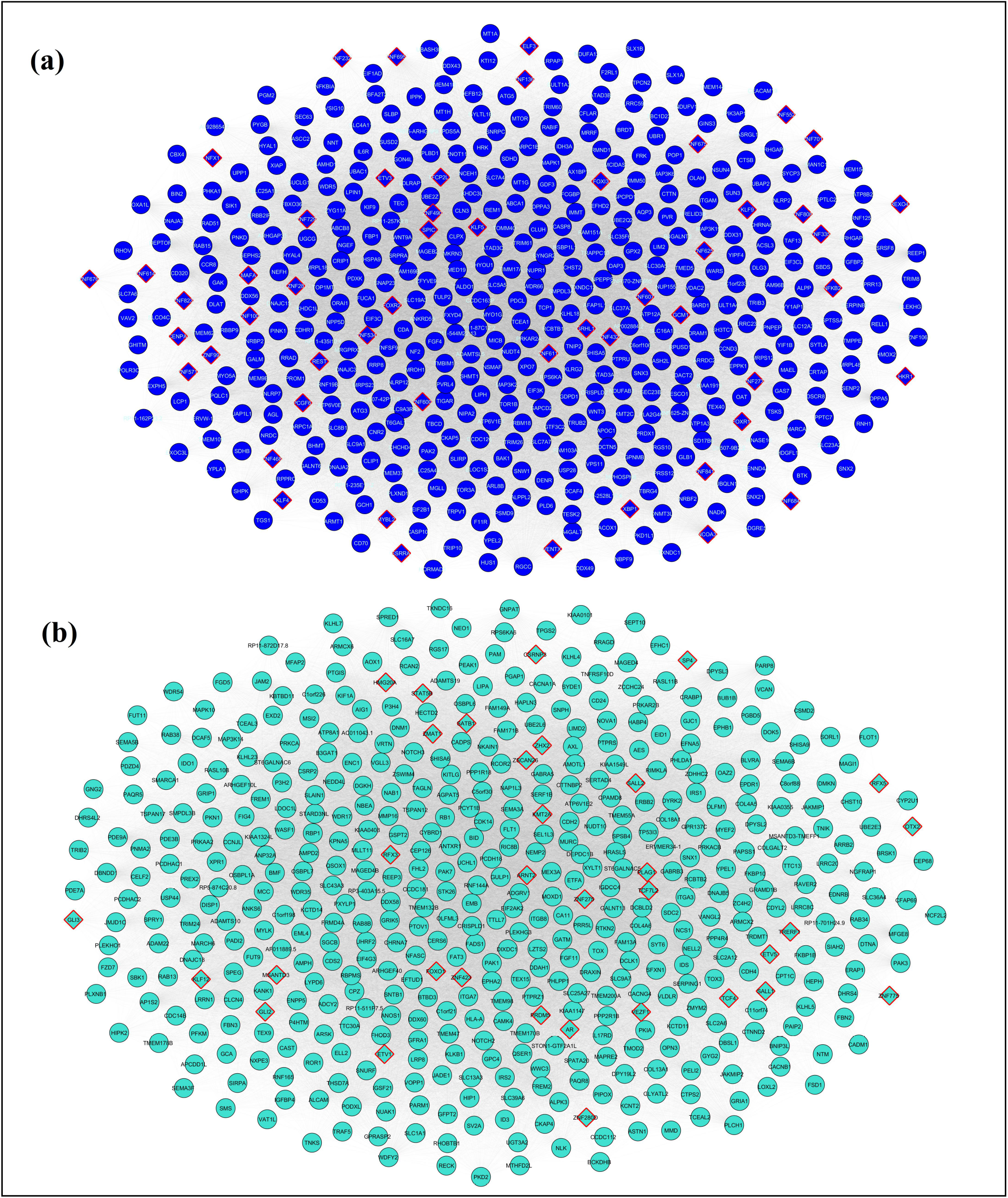
Network of hub genes in (a) blue and (b) turquoise modules connected by the top 1% edges by weight within each module. The blue module hub gene network contains 479 nodes connected by 41,338 edges while the turquoise module hub gene network contains 507 nodes and 43,171edges. The diamond-shaped nodes here represent the transcription factors. An interactive Cytoscape file of the networks is provided as Supplementary File 5.

### 3.5. Analysis of hub gene expression pattern

Further, we wanted to see if the hub genes showed significant changes in their gene expression between the two different states. Differential gene expression analysis was performed and the genes preferentially expressed in naive (|log2fc| < −1.0 and p_adj_ < 0.05) and primed (|log2fc| >1.0 and p_adj_ < 0.05) states identified. We observed, 448 of 479 blue module hub genes were indeed preferentially expressed in the naive state whereas 503 of 507 turquoise module hub genes were preferentially expressed in the primed state (Figure 8a). This comes in support of the gene expression heatmap of the module genes (Figure 5) which showed the blue module to contain naive specific genes and turquoise module contain primed specific genes. However, when we assessed the overlap of the blue and turquoise module hub genes with the top 479 and 507 (by log_2_fold-change) naive and primed state preferred genes, we observed 213 and 252 genes overlap respectively (Figure 8b). Thus, this shows that though the intra-modular hub genes are significantly differentially expressed, they need not have high fold-change. Alternately, this reaffirms the fact that differential gene expression analysis alone is not sufficient for identifying the critical genes involved in a biological process and must be combined or preceded by network-based approach [45]. The intra-modular hub genes in the blue and turquoise modules were found to have an average log_2_FoldChange of 2.71 and 3.13 respectively (Figure 8c).

**Figure 8:**
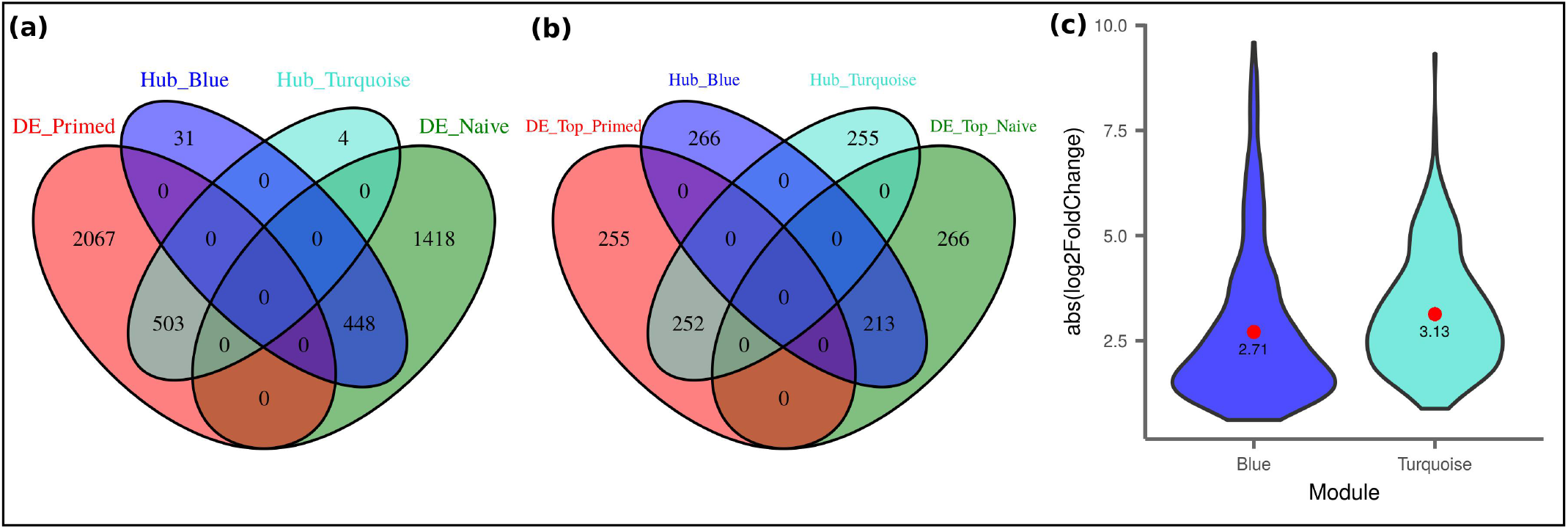
(a) Venn diagram showing the overlap between the intra-modular hub genes and the genes preferentially expressed in the two pluripotency states. (b) Venn diagram showing the overlap of the intra–modular hub genes with the top 479 and 507 (by log2FoldChange) naive and primed state preferred genes. (c) Violin plot representing the log2FoldChanges of the intra-modular hub genes. The value within the violin indicated the average log2FoldChange values of the hub genes.

### 3.6. Transcription factor screening among intra-modular hub genes

Transcription factors (TFs) are key elements that regulate gene expression. They have been known to play a crucial role in cell differentiation and de-differentiation. In order to identify the TFs among the blue and turquoise module hub genes, we compared them with a manually curated list of 1639 human TFs by

Lambert et al [46]. We found 52 and 33 TFs among the blue and turquoise module hubs respectively (Figure 9, Supplementary File 6). Interestingly, we observed that at least 50% of the TFs in each group were zinc finger proteins (ZNFs). In particular, the C2H2-type ZNFs were most abundant (32 in blue and 16 in turquoise modules). The importance of zinc finger proteins in pluripotency and human development is well established. For example, the Krüppel-like factor family members KLF4 and KLF5 along with KLF2 has been found to be sufficient to support self-renewal and their removal can compromise it [47]. KLF4 and another blue module hub transcription factor TFCP2L1 are a part of the core gene regulatory network of naive pluripotency in mESC [48]. ZNF534 is a negative regulator of the human endogenous retrovirus HERVH which in turn has a key regulatory role in human PSCs [17]. SALL1 and SALL2 are key regulators of organogenesis [49].

**Figure 9:**
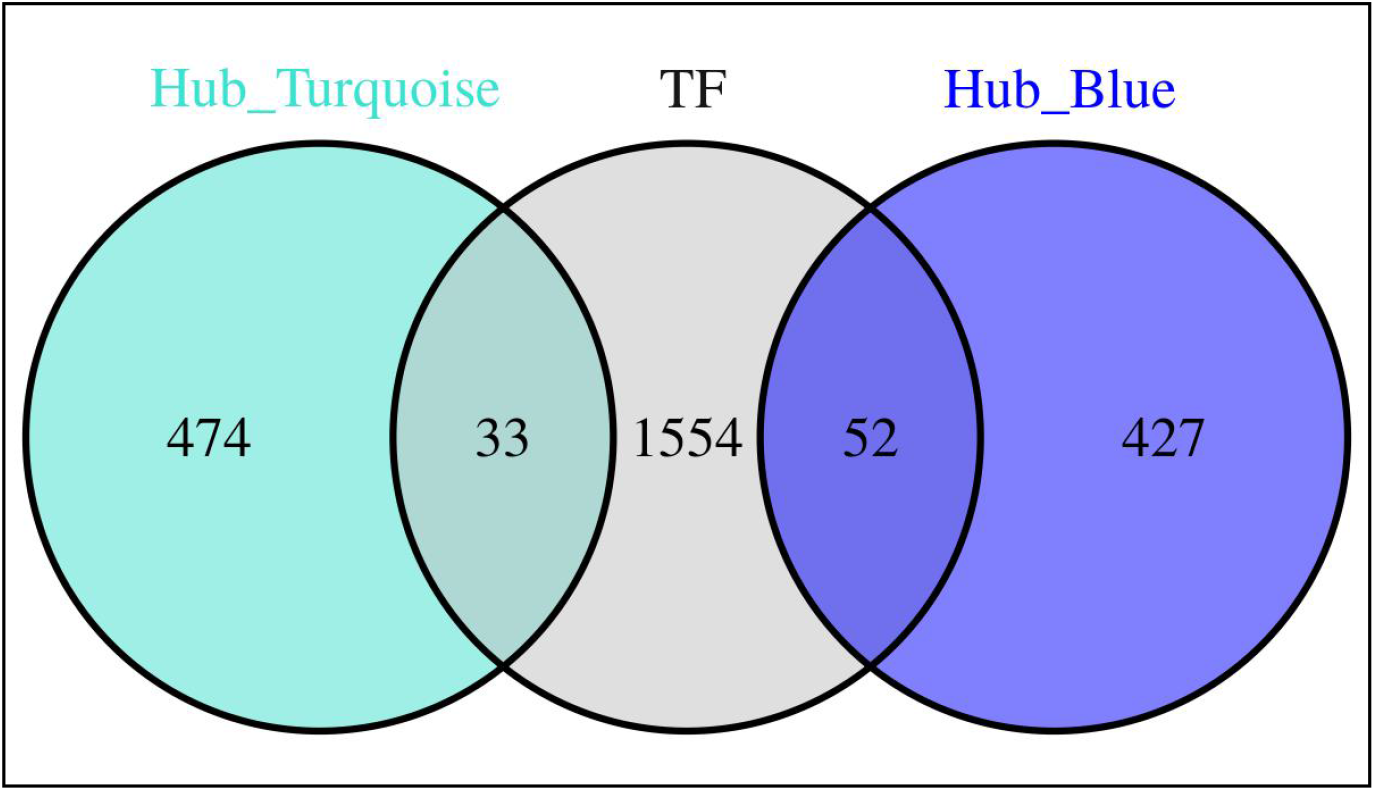
Venn diagram depicting the overlap of the intra–modular hub genes with the manually curated transcription factor list.

Besides the ZNFs, other blue module hub TFs include MYBL2 which regulates cell cycle progression and is essential for mouse inner cell mass formation [50], ESRRA a key regulator of mitochondrial biogenesis and energy homeostasis [51], REST that has been found to promote maintenance of neural stem cells and prevent premature differentiation into mature neurons [52], and NCOA3 which has been reported to control Esrrb-dependent activation of important self-renewal regulators including KLF4 in mESCs [53]. Interestingly here though, ESRRB does not occur in any of our gene lists and it has been reported that ESRRB is expressed in the mouse epiblast but not in the human epiblast [54]. Among the turquoise module hub TFs include OTX2 which is one of the first TFs which is activated during exit from the naive state of pluripotency [55] and FOXO1 an essential regulator of pluripotency in hESCs [56]. The occurrence of these TFs as hubs in the networks of naive and primed pluripotent states suggest their role as master regulators behind the two pluripotent stem cell states. Further, as these TFs are specific to either of the two conditions, they might play a switch on-off mechanism in forming and maintaining the two pluripotent states (Figure 10). This is in consistency with previous reports that the transcriptional network required to maintain the expression of core TFs, which is completely distinct in the two pluripotent states [3,47].

**Figure 10:**
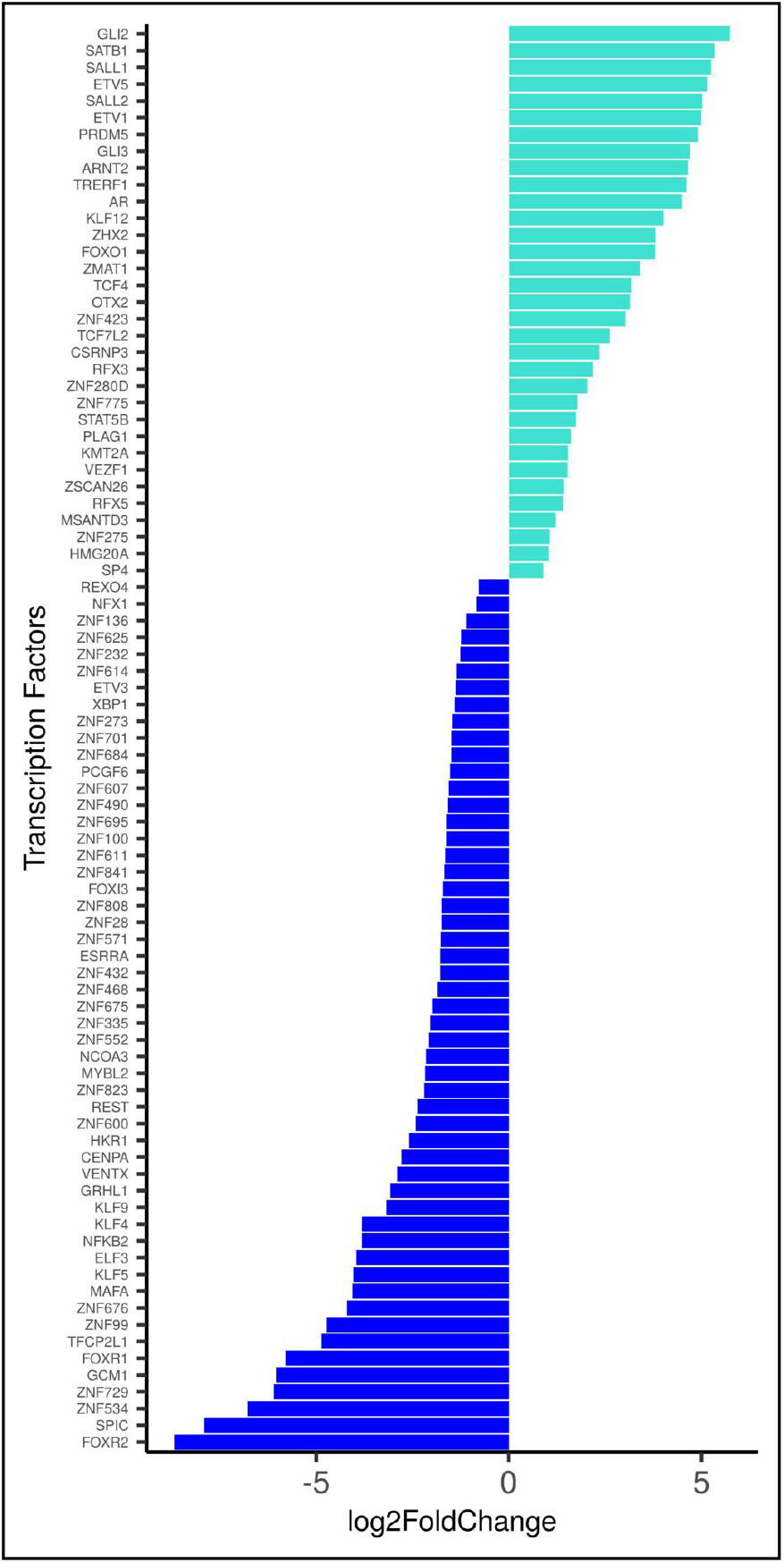
Bar–plot depicting the fold-change (on log2 scale) of the transcription factor hubs. The colour of the bars indicates the module to which they belong.

## 4. CONCLUSIONS

The naive and primed states of pluripotency represent the early stages of human development. In this work, we used a WGCNA approach for identifying the key genes that differentiate the two distinct pluripotent states in human. The identification of these molecular markers defining the two states would aid in better understanding of the process of human development and ultimately in biomedical applications. In all, we identified 4791 genes (blue module) to be involved in naive state and 5066 genes (turquoise module) to be involved with the primed state. The involvement of the genes in the two states were verified by functional and pathway enrichment analysis. We further refined this list and identified 479 hub genes for naive and 507 hub genes for primed states. These state-specific hubs play key roles in the development timeline of the naive and primed states. We also identified the key TFs that might be the master regulators in formation of the two pluripotency states. In fact, gene expression patterns of these TFs suggests a somewhat switch on-off mechanism that guides naive and primed states of pluripotency in human. Thus, our work generates an important resource of transcriptomic markers in naive and primed PSCs and provides a framework for the future investigation of human pluripotency. We strongly believe these reported genes would guide the experimentalists in designing downstream experiments and will ultimately help to understanding the underlying regulatory network defining pluripotency.

## CONFLICTS OF INTEREST

The authors declare that they have no conflict of interest.

## ACKNOWLEDGEMENTS

AS thank the Department of Biotechnology (DBT) for providing financial assistance for the project. Thanks to the members of Som Lab (https://www.somlab.in/) for their helpful discussions. AS acknowledges useful discussions with Wentian Li.

## FUNDING

The work was supported by funding from the Department of Biotechnology (DBT), Government of India (Grant No. BT/PR12842/BID/7/521/2015)

## ABBREVIATIONS

CPM: Count Per Million
ENA: European Nucleotide Archive
FDR: False Discovery Rate
GO: Gene Ontology
hESC: Human Embryonic Stem Cell
mESC: Mouse Embryonic Stem Cell
PSC: Pluripotent Stem Cell
TF: Transcription Factor
TOM: Topological Overlap Measure
WGCNA: Weighted Gene Co-expression Network Analysis
ZNF: Zinc Finger Protein

## SUPPLEMENTARY FILES

**Supplementary File 1:** Details of raw RNA–seq data included in the study.

**Supplementary File 2:** List of genes in the blue (naive) and turquoise (primed) modules.

**Supplementary File 3:** Results of functional and pathway enrichment analysis.

**Supplementary File 4:** List of hub genes in the blue (naive) and turquoise (primed) modules.

**Supplementary File 5:** Cytoscape file of the networks.

**Supplementary File 6:** List of transcription factors identified as hubs in the blue (naive) and turquoise (primed) modules.

## Notes

### Competing Interest Statement

The authors have declared no competing interest.

